# Disease-associated *MYH11* tail variants disrupt vascular smooth muscle myosin II filament dynamics and cell mechanics

**DOI:** 10.64898/2026.10.22.760788

**Authors:** Sasha K. Demeulenaere, Margaret A. Bennett, Hiral Patel, Gregory Aubert, Dianna M. Milewicz, Patrick W. Oakes, Jordan R. Beach

## Abstract

Smooth muscle myosin II (SMII), encoded by *MYH11*, is the principal motor driving vascular smooth muscle cell (VSMC) contraction. How disease-associated *MYH11* variants alter SMII behavior in living cells remains poorly understood. We directly visualized two human *MYH11* tail variants in VSMCs: pathogenic K1256del and a variant of uncertain significance A1839V, identified in a patient with severe aortoiliac aneurysmal disease. Both variants reduced SMII filament assembly and VSMC contractile output, but with distinct severity. A1839V predominantly retained filamentous organization and displayed partial agonist-induced stabilization, consistent with an intermediate cellular phenotype. In contrast, K1256del severely disrupted filament architecture and behavior, including reduced focal adhesion organization, increased SMII network flow and accelerated filament exchange. Furthermore, K1256del abolished the filament stabilization response to induced contraction. These findings identify failure of SMII filament stabilization as a cellular mechanism linking *MYH11* variation to impaired VSMC mechanics. More broadly, they show that direct live-cell phenotyping can distinguish variant-specific defects in the contractile machinery, providing a scalable functional strategy to complement genetic classification and prioritize *MYH11* variants for deeper mechanistic and in vivo study.

**Significance Statement:** Thoracic aortic aneurysms and dissections can arise when the muscular wall of the aorta is weakened and cannot withstand the pressure of blood flow. Mutations in *MYH11*, which encodes smooth muscle myosin II, are linked to inherited aortic disease, but how these variants alter vascular smooth muscle cell function remains unclear. Here, we directly visualize two *MYH11* variants in living cells and measure how they disrupt myosin organization, stability, and force generation to different degrees. The pathogenic K1256del variant causes severe defects and prevents myosin stabilization during contraction, whereas the A1839V variant of uncertain significance causes milder defects. These findings reveal how *MYH11* variants impair vascular smooth muscle function and provide a cell-based approach for evaluating poorly understood genetic variants.

## Introduction

Vascular smooth muscle cells (VSMCs) form the medial layer of arteries and provide the contractile force required for vascular tone and vessel wall integrity [1, 2]. In large arteries such as the aorta, VSMCs are organized within elastin lamellar units and mechanically coupled to elastin and other extracellular matrix proteins, allowing cellular contraction to contribute to tissue-level mechanics and resistance to pulsatile pressure [3, 4, 5]. Defects in VSMC contraction or force transmission contribute to aneurysm formation and dissections in the vessel wall [6, 7].

VSMC force generation is driven by the actomyosin cytoskeleton, with smooth muscle myosin II (SMII) serving as the principal motor [8, 7]. SMII monomers assemble into bipolar filaments through coiled-coil heavy-chain tail electrostatic interactions, and drive contraction using motor domains to bind and pull on the actin thin filaments [9, 10, 11]. This contractile force is transmitted through actomyosin fibers and focal adhesion networks to the surrounding matrix [12, 13]. Thus, SMII is not only a molecular motor, but also part of a larger mechanical system that links SMII filament assembly dynamics to cell-scale force generation and vessel wall stability.

Our previous work showed that SMII filaments are not static contractile structures [14]. At baseline, SMII is predominantly filamentous but remains highly dynamic, exchanging between filament and monomer pools. During agonist-induced contraction, SMII further assembles and stabilizes, suggesting that VSMC contractility depends not only on activating pre-existing myosin motors, but also on regulating SMII filament assembly and exchange [14]. This dynamic assembly-stabilization cycle provides a framework for understanding how defects in SMII filament regulation could impair VSMC mechanics. Because VSMC force is transmitted through cytoskeletal, focal adhesion, and elastic lamina connections, defects in SMII assembly could propagate from molecular filament instability to tissue-level wall failure [5, 4, 15, 7].

The SMII heavy chain is encoded by *MYH11*, and pathogenic variants in the gene provide a direct genetic link between the contractile machinery and inherited thoracic aortic disease [16, 17, 18, 7]. *MYH11* variants were first identified in families with thoracic aortic aneurysm and dissection (TAAD) with patent ductus arteriosus (PDA) [16]. Subsequent work has shown that *MYH11*-associated disease is characterized by abnormal smooth muscle organization, altered vessel wall mechanics, and impaired contractile function [17, 5, 19, 20]. The pathogenic *MYH11* K1256del variant was identified in families with heritable thoracic aortic disease and later modeled in mice, where it reduced aortic contractility and vessel wall structural integrity and increased susceptibility to aortic dissection [21, 20]. However, even for this well-studied variant, it remains unclear how the mutation alters SMII filament organization and dynamics within individual VSMCs. This places SMII well behind its striated and non-muscle myosin 2 (NM2) paralogs, for which disease-associated mutations have been studied extensively and myosin-directed therapies have advanced substantially, including the development of mavacamten for hypertrophic cardiomyopathy and selective inhibitors targeting NM2 [22, 23, 24, 25]. Additionally, understanding how *MYH11* pathogenic variants disrupt SMII filaments is especially relevant for variants of uncertain significance (VUS), which are increasingly identified by clinical genetic testing and often require functional data to determine their biological significance [26]. A scalable cell-based approach that directly visualizes variant SMII behavior could help bridge the gap between sequence-based variant identification and whole-animal disease modeling.

In this work, we analyzed two *MYH11* variants in the tail of SMII in human VSMCs: the pathogenic *MYH11* K1256del variant and the novel VUS *MYH11* A1839V. We characterized their effects on SMII filament organization, cytoskeletal and focal adhesion architecture, contractile force generation, SMII assembly and exchange dynamics, and agonist-induced filament stabilization. To our knowledge, this is the first study to directly visualize these *MYH11* tail variants in living VSMCs. Both variants altered SMII behavior, but with distinct degrees of severity. A1839V produced measurable defects in SMII assembly and cell mechanics despite its uncertain clinical significance, whereas K1256del produced a more severe phenotype characterized by disrupted filament architecture, reduced force generation, impaired focal adhesion organization, increased network flow, accelerated SMII exchange, and failure to stabilize in response to contractile activation. Together, these findings establish direct cellular phenotyping as a successful strategy to define how individual *MYH11* variants alter the SMII cytoskeletal machinery and to prioritize variants for deeper mechanistic or in vivo study.

## Results

### *MYH11* tail variants alter SMII filament architecture in VSMCs.

We first sought to determine whether *MYH11* tail variants directly alter SMII organization in VSMCs. We focused on two variants with different levels of clinical evidence: K1256del, a pathogenic *MYH11* tail variant previously linked to heritable thoracic aortic disease [21], and *MYH11* A1839V, a VUS identified in a Loyola Medicine patient with severe infrarenal abdominal aortic and bilateral common iliac aneurysmal disease requiring repeated endovascular repair but not enlargement of the proximal aorta. The patient also had a family history notable for abdominal aortic aneurysm in multiple first-degree relatives but no history of PDAs or thoracic aortic disease. A1839V has appeared in public variant databases and in a supplemental genomic sequencing dataset, but available reports do not establish its functional consequence in VSMCs or its role in aortic disease [27]. A1839V is also listed in ClinVar with conflicting interpretations, including uncertain significance and likely benign classifications, with submitter comments noting limited supporting evidence and no reported functional studies [28]. Like many VUS, these conflicting data points make A1839V a poor candidate for financially- and temporally-demanding animal studies, but ideal test-case VUS to explore with more molecular precision in cell-scale studies.

To characterize wild-type and *MYH11* tail variants in human primary VSMCs, we generated adenoviral EGFP-SMII constructs with EGFP fused to the N-terminus of the SMII heavy chain (Fig. 1A). Western blot analysis confirmed expression of wild-type, A1839V, and K1256del EGFP-SMII above endogenous SMII, as expected, with EGFP signal detected only in cells expressing tagged SMII constructs (Fig. 1B). The exogenous EGFP-SMII to endogenous SMII ratio was high for all three conditions, suggesting relatively minimal contribution of the endogenous SMII (Fig. 1C). We then asked whether these variants altered SMII localization and filament architecture. Wild-type EGFP-SMII primarily incorporated into long actin-associated stress fibers and dense network-like filament arrays (Fig. 1D). A1839V retained filamentous organization but showed a shift toward shorter, more disorganized stress fiber-like structures. In contrast, K1256del produced a more severe disruption of SMII morphology, with few long stress fibers and more frequent cytoplasmic and punctate aggregate patterns (Fig. 1D,E). Additionally, K1256del expressing cells displayed reduced actin stress fiber organization.

**Figure 1.**
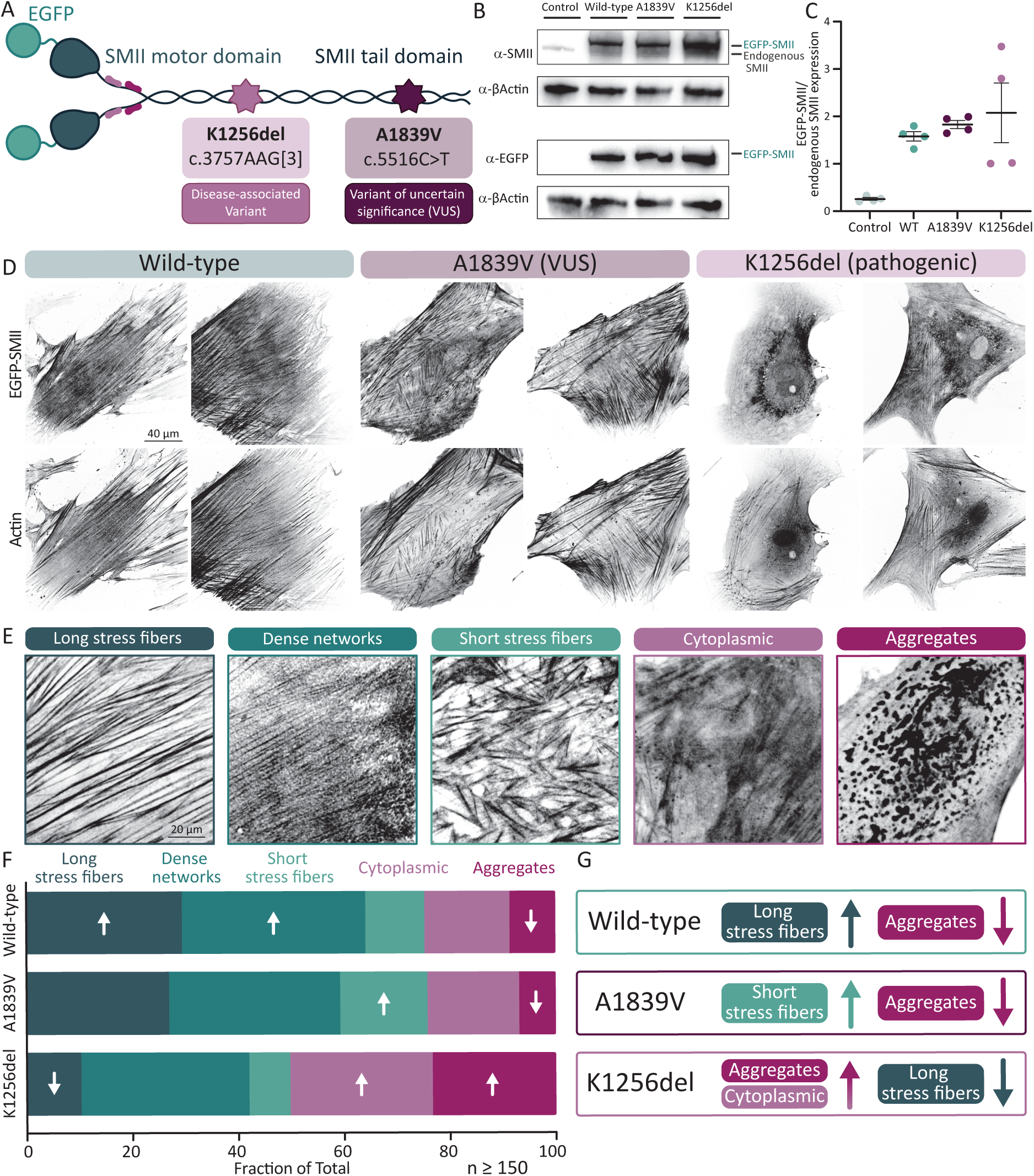
*MYH11 tail variants alter SMII filament architecture in VSMCs.* **A)** Schematic of N-terminally EGFP-tagged SMII constructs, with the locations of the A1839V and K1256del variants indicated. **B)** Western blots of control VSMCs and VSMCs expressing wild-type, A1839V, or K1256del EGFP-SMII probed for SMII, EGFP, and *β* -actin. **C)** Quantification of EGFP-SMII expression relative to endogenous SMII. Each dot represents a separate viral transduction. **D)** Duplicate representative EGFP-SMII and phalloidin-stained actin images from human VSMCs expressing wild-type, A1839V, or K1256del EGFP-SMII. **E)** Representative examples of the five SMII morphology categories: long stress fibers, dense networks, short stress fibers, cytoplasmic, and aggregates. **F)** Scorer-averaged fractions of morphology classifications from three blinded reviewers (*≥* 150 cells per variant). Overall morphology distributions differed across variants by chi-square test of independence on a scorer-normalized 3 *×* 5 contingency table (*χ*^2^ (8)= 107.24, *p <* 0.0001). Upward and downward arrows indicate variant-morphology combinations observed more or less frequently than expected, respectively, among combinations with absolute Pearson residuals *>* 2; residual values are reported in Table S1. **G)** Summary of the morphology categories most statistically enriched or depleted relative to expected distributions for each variant

To quantify these changes, we performed blind classification of EGFP-SMII architecture into five reproducible categories: long stress fibers, dense networks, short stress fibers, cytoplasmic, and aggregates, where individual cells could be classified into multiple categories (Fig. 1E). Architecture distributions differed significantly across wild-type, A1839V, and K1256del EGFP-SMII (*χ*^2^(8)= 107.24, *p <* 0.0001; Fig. 1F,G; Table S1). Wild-type was enriched for long stress fibers and depleted for aggregates, consistent with robust assembly of EGFP-SMII into organized filament networks. *MYH11* A1839V was enriched for short stress fibers and depleted for aggregates (Fig. 1F,G; Table S1), suggesting a milder disruption that preserves filamentous organization but alters network architecture. *MYH11* K1256del was the strongest driver of the altered distribution, with enrichment of cytoplasmic and aggregate morphologies and marked depletion of long stress fibers (Fig. 1F,G; Table S1). Together, these data show that *MYH11* tail variants produce distinct defects in higher-order SMII organization, with pathogenic K1256del causing the most severe loss of normal filament architecture.

While we observed distinct phenotype distributions between wild-type, *MYH11* A1839V, and K1256del, the architectures were heterogeneous within each group. For example, we observed a minority of VSMCs expressing K1256del with apparently preserved SMII architecture (Fig. S1A). This heterogeneity raised an important mechanistic question: do *MYH11* tail variants produce structurally abnormal filaments when they incorporate into stress fibers, or do they primarily reduce the frequency or stability of normal filament assembly? To identify individual filamentous units, we performed super-resolution SIM imaging on cells expressing wild-type, *MYH11* A1839V, and K1256del EGFP-SMII that were co-stained for NM2A, which we previously demonstrated could co-assemble with SMII [14]. Because EGFP marks the N-terminus of SMII and the NM2A antibody labels the C-terminus of endogenous NM2A (Fig. S1B), filaments appear as NM2A tail puncta flanked on either side by EGFP-SMII head puncta. We observed extensive overlap between SMII and NM2A within filamentous structures throughout the cell (Fig. S1C,E,G). Closer examination of individual fibers revealed distinct head-tail-head alternating patterns of the two fluorophores (Fig. S1D,F,H), demonstrating that SMII and NM2A are co-organized within the same contractile filament bundles, as has been seen previously for SMII and NM2 isoforms [29, 30, 14]. The alternating SMII-NM2A-SMII (green-magenta-green) patterns are also *∼*300 nm in length, consistent with what would be seen in co-assembled bipolar filaments. We observed this alternating pattern in all SMII variants when stress fibers were present. We also found that the aggregates seen in the K1256del VSMCs contained both SMII and NM2A, suggesting that K1256del does not simply exclude NM2A from SMII-containing structures, but instead may disrupt the normal organization of mixed myosin 2 assemblies (Fig. S1I,J). These findings demonstrate that potential and confirmed pathogenic SMII variants are capable of forming relatively normal filaments and architectures, but do so with a significantly reduced frequency and capacity.

### *MYH11* tail variants reduce VSMC contractile force generation.

Because wild-type and variant EGFP-SMII displayed distinct filament morphologies, we hypothesized these structural differences would disrupt force transmission through the actomyosin-adhesion system. Productive VSMC force generation requires SMII filaments to couple to actin stress fibers and transmit tension through focal adhesions to the extracellular matrix [13, 12]. We therefore examined phospho-paxillin, a marker of force-associated focal adhesions [31]. Wild-type cells displayed phospho-paxillin-positive adhesions along the cell periphery and at the ends of organized SMII-containing stress fibers, and A1839V cells retained qualitatively similar adhesion organization in regions with preserved filamentous SMII (Fig. 2A). In contrast, K1256del cells showed less apparent coordination between SMII structures and phospho-paxillin-positive adhesions. Quantification of phospho-paxillin-positive focal adhesion area normalized to cell area showed a significant reduction in K1256del cells compared with wild-type (Fig. 2B,C), consistent with impaired maturation or maintenance of force-associated adhesions. To directly test how expression of the SMII variants impacts cell contractility, we measured their force production using traction force microscopy. Cells expressing wild-type EGFP-SMII generated robust traction stresses (Fig. 2D). In contrast, cells expressing A1839V or K1256del EGFP-SMII exhibited significantly reduced overall force generation, quantified as contractile energy per area (Fig. 2E). These differences were not explained by exogenous EGFP-SMII expression level, as the contractile work done by the cell did not show an obvious relationship with EGFP-SMII fluorescence intensity across groups (Fig. S2A). We also noted that expression of both the *MYH11* A1839V and K1256del variants resulted in a shift in the distribution of forces away from the center of the cell towards the periphery, consistent with the reduction in focal adhesions and reduced stress fiber formation (Fig. S3). Together, these data show that *MYH11* tail variants disrupt VSMC mechanics at two linked levels: impaired organization of the adhesion structures that transmit actomyosin tension to the extracellular matrix and reduced cellular-scale contractile output.

**Figure 2.**
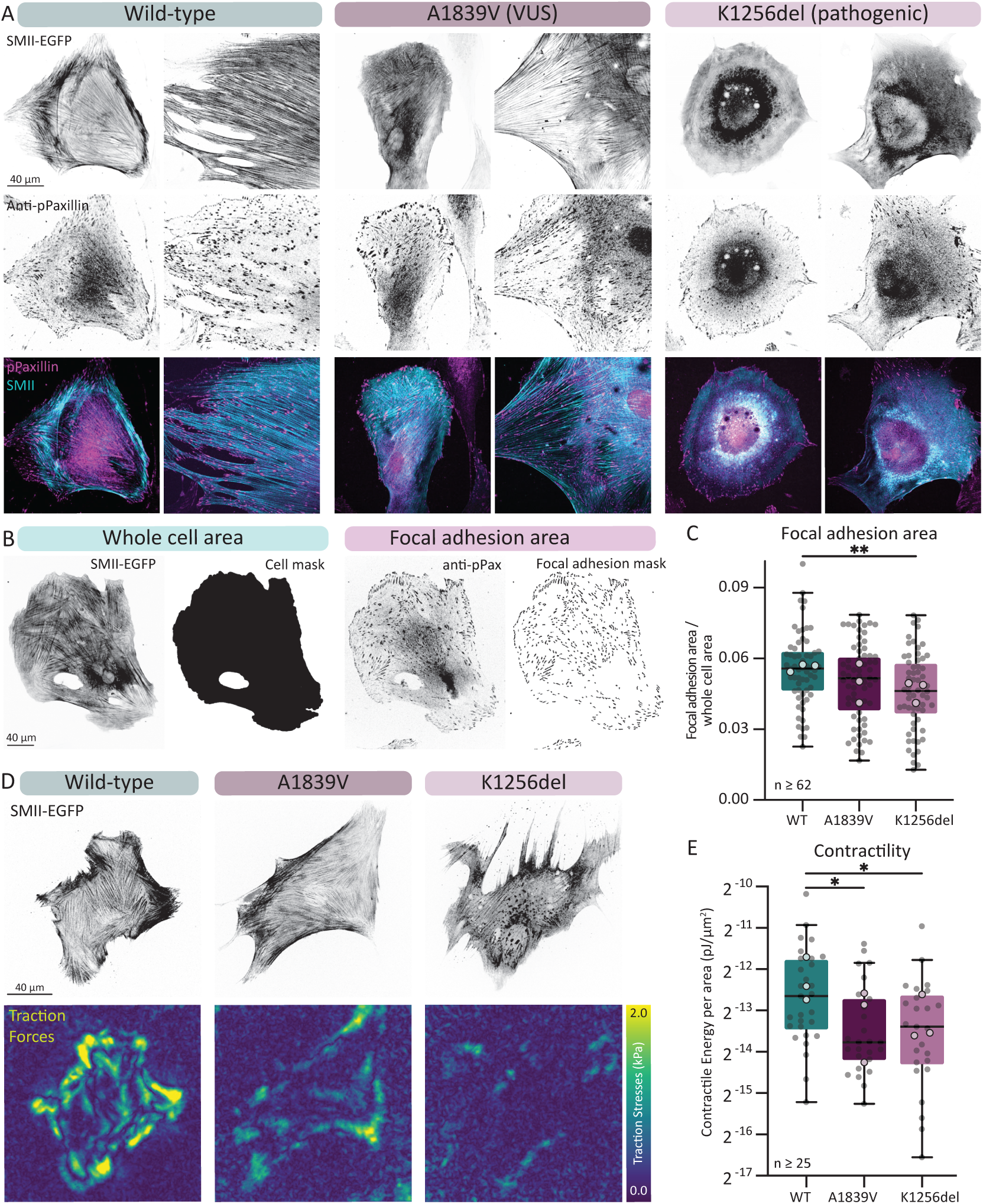
*MYH11 tail variants alter adhesion organization and reduce VSMC force generation.* **A)** Duplicate representative images of VSMCs expressing wild-type, A1839V, or K1256del EGFP-SMII and immunostained for the focal adhesion protein phospho-paxillin. **B)** Representative whole-cell and focal adhesion masks used to quantify phospho-paxillin-positive adhesion area normalized to whole-cell area. **C)** Quantification of focal adhesion area normalized to whole-cell area in cells expressing wild-type, A1839V, or K1256del EGFP-SMII. **D)** Representative images of VSMCs expressing wild-type, *MYH11* A1839V, or K1256del EGFP-SMII and corresponding traction stress maps generated by traction force microscopy. **E)** Quantification of contractile energy per area for cells expressing wild-type, A1839V, or K1256del EGFP-SMII. Data shown in C and E are displayed as individual cells with box-and-whisker plots showing the median and interquartile range (C, *≥* 62 cells per group; E, *≥* 25 cells per group). Small gray points represent individual cells, and larger points represent the mean value for each independent experimental replicate. Groups were compared by Kruskal-Wallis test followed by Dunn’s multiple comparisons test because the distributions were not normally distributed. * indicates *p <* 0.05; ** indicates *p <* 0.01

### K1256del increases SMII filament network flow, decreases SMII assembly, and accelerates SMII exchange kinetics.

The reduced adhesion organization and force generation observed in variant-expressing cells suggested that *MYH11* tail variants might alter not only the steady-state architecture of SMII, but also network dynamics. To quantify network dynamics directly, we acquired time-lapse images of EGFP-SMII expressing VSMCs and used dense optical flow analysis to measure relative motion in the cytoskeletal network (Fig. 3A; Movie M1). Flow maps depict the local magnitude and direction of SMII network movement, whereas cell-level mean flow was calculated by averaging flow-vector magnitudes across the cell area and imaging period. Wild-type EGFP-SMII networks showed relatively coherent, low-amplitude flow within organized filament arrays, consistent with a stable contractile network. A1839V networks did not differ significantly from wild-type networks, but did show increased data point variability in mean network flow (Fig. 3B). In contrast, K1256del-expressing cells displayed increased mean network flow across disorganized SMII structures (Fig. 3B). Mean network flow was not associated with EGFP-SMII fluorescence intensity, indicating that the K1256del phenotype was not explained by differences in exogenous SMII expression (Fig. S2B). Rather, it is consistent with a less stable and more dynamic SMII filament network.

**Figure 3.**
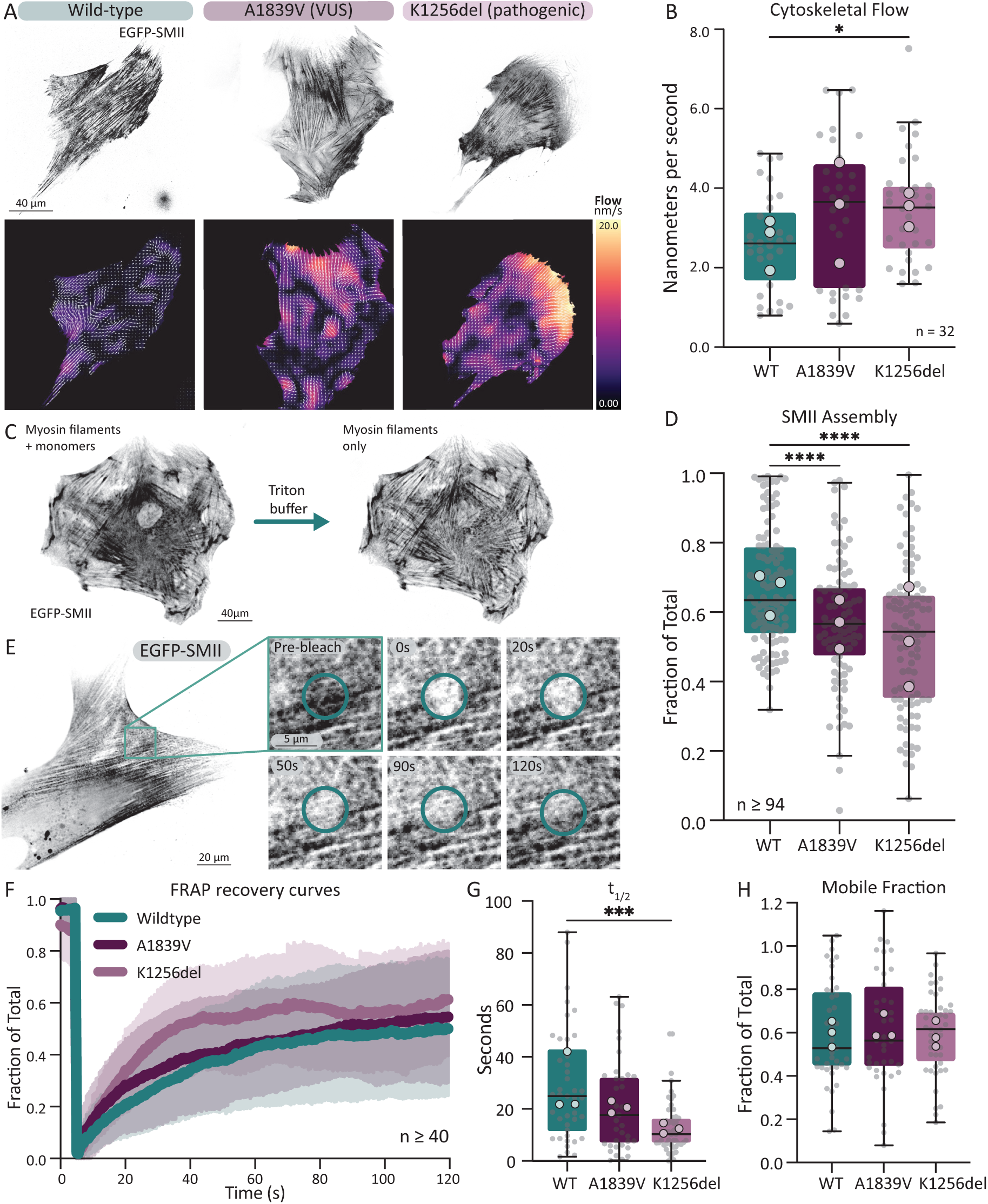
*K1256del increases SMII filament network flow, decreases SMII assembly, and accelerates SMII exchange kinetics.* **A)** Representative EGFP-SMII images and dense optical flow maps from VSMCs expressing wild-type, A1839V, or K1256del EGFP-SMII. Flow maps indicate local network direction and velocity. See Movie M1. **B)** Mean SMII network flow, calculated by averaging flow-vector magnitudes across the cell area and imaging period (*n* = 32 cells per group). Each variant was compared with wild-type using pairwise Mann-Whitney U tests. * indicates *p <* 0.05. **C)** Representative images from a Triton permeabilization assay showing total EGFP-SMII before permeabilization and the remaining filament-associated pool after removal of soluble SMII. **D)** Quantification of the fraction of EGFP-SMII assembled (*≥* 94 cells per group) as measured in a Triton permeabilization assay. **E)** Representative images from a FRAP experiment showing recovery of EGFP-SMII in a bleached region during recovery. **F)** Mean FRAP recovery curves for wild-type, A1839V, and K1256del EGFP-SMII. Recovery curves are shown as mean *±* SEM and were fit to a single exponential. **G,H)** Quantification of G) *t*_1_*_/_*_2_ and H) mobile fraction (*≥* 40 cells per group). Recovery curves were fit to a single exponential. For assembly and FRAP analyses, each variant was compared with wild-type using a separate unpaired t test. For B, D, G and H, small gray points represent individual cells and larger points represent independent experimental-replicate means. *** indicates *p <* 0.001; **** indicates *p <* 0.0001.

We next asked whether this increased network flow reflected altered steady-state partitioning of SMII between filament-associated and soluble pools. We performed a single-cell imaging-based assembly assay to measure the fraction of SMII assembled into filaments [14]. VSMCs expressing wild-type, A1839V, or K1256del EGFP-SMII were first imaged to measure total EGFP-SMII intensity and then permeabilized with a physiological buffer containing Triton to extract the soluble SMII pool. The remaining Triton-insoluble EGFP-SMII signal represented the filament-associated fraction (Fig. 3C). Wild-type EGFP-SMII was predominantly filament-associated at steady state (Fig. 3D). In contrast, both A1839V and K1256del displayed heterogeneous assembly levels with significantly reduced fractions of assembled EGFP-SMII (Fig. 3D).

To determine whether the variants also altered SMII filament exchange kinetics we used Fluorescence Recovery After Photobleaching (FRAP). A small region of the EGFP-SMII network was photobleached, and fluorescence recovery was monitored to quantify the fraction of SMII participating in exchange (mobile fraction) and the time it takes for half of the mobile fraction to exchange (*t*_1_*_/_*_2_; Fig. 3E & F). While A1839V did not significantly alter *t*_1_*_/_*_2_, it showed a trend towards increased exchange kinetics (*p* = 0.18). K1256del significantly decreased *t*_1_*_/_*_2_ compared with wild-type, indicating faster exchange of SMII within the filament network (Fig. 3G). Neither variant significantly changed the mobile fraction, indicating that the overall fraction of the exchanging SMII pool remained similar across groups (Fig. 3H).

Combined, these results demonstrate that A1839V and K1256del both reduce steady-state SMII filament assembly but have distinct impacts on filament exchange kinetics. A1839V reduced the fraction of assembled SMII while having minimal impact, if any, on exchange and network flow, whereas K1256del both reduced assembly and accelerated exchange and network flow. Thus, K1256del produces a less stable, more dynamically exchanging SMII filament network, while A1839V produces an intermediate phenotype, reducing assembly without producing the same clear increase in network flow or exchange kinetics.

### K1256del abolishes agonist-induced stabilization of SMII filaments.

In our previous work, we found that while SMII is predominantly filament-associated, it remains highly dynamic at baseline. Agonist stimulation, however, increased SMII filament assembly while reducing exchange in cultured VSMCs, primary aortic VSMCs, and intact arterioles, demonstrating agonist-induced stabilization of SMII filaments [14]. These findings suggested that physiological VSMC contraction depends not only on activating pre-existing SMII filaments, but also on dynamically shifting SMII into a more stable filament-associated state. We therefore asked whether *MYH11* tail variants retained the ability to stabilize in response to contractile activation.

Carbachol is a cholinergic receptor agonist that stimulates IP_3_-dependent release of intracellular calcium in VSMCs [32], leading to increased myosin regulatory light chain phosphorylation by myosin light chain kinase, which in turn promotes SMII activation and contraction. To test whether agonist stimulation stabilizes the two variants, we first performed FRAP in one region of a VSMC under control conditions, added carbachol, and then rapidly performed FRAP in a second, distinct region of the same cell for comparison (Fig. 4A). This paired approach provided a cell-specific internal control for measuring agonist-induced changes in SMII exchange kinetics (Fig. 4A). As expected, wild-type EGFP-SMII exhibited a robust stabilization response following carbachol treatment, with a decreased mobile fraction and increased *t*_1_*_/_*_2_ indicating a reduction in the size of the exchanging SMII pool and slower exchange of SMII within filamentous structures (Fig. 4B). A1839V filaments retained part of this agonist-induced response. Carbachol significantly decreased the mobile fraction of A1839V EGFP-SMII, although to a lesser extent compared to wild-type, indicating that the variant remained capable of filament stabilization following stimulation (Fig. 4C). However, carbachol did not significantly alter A1839V *t*_1_*_/_*_2_. In contrast, K1256del failed to respond to carbachol. Neither the mobile fraction nor *t*_1_*_/_*_2_ changed significantly following stimulation, indicating a loss of normal agonist-induced regulation of SMII exchange kinetics (Fig. 4D). This absence of a stabilization response represents a major functional defect. Although wild-type SMII shifts into a more stable filament state during contractile activation, K1256del remains dynamically unresponsive to the same stimulus.

**Figure 4.**
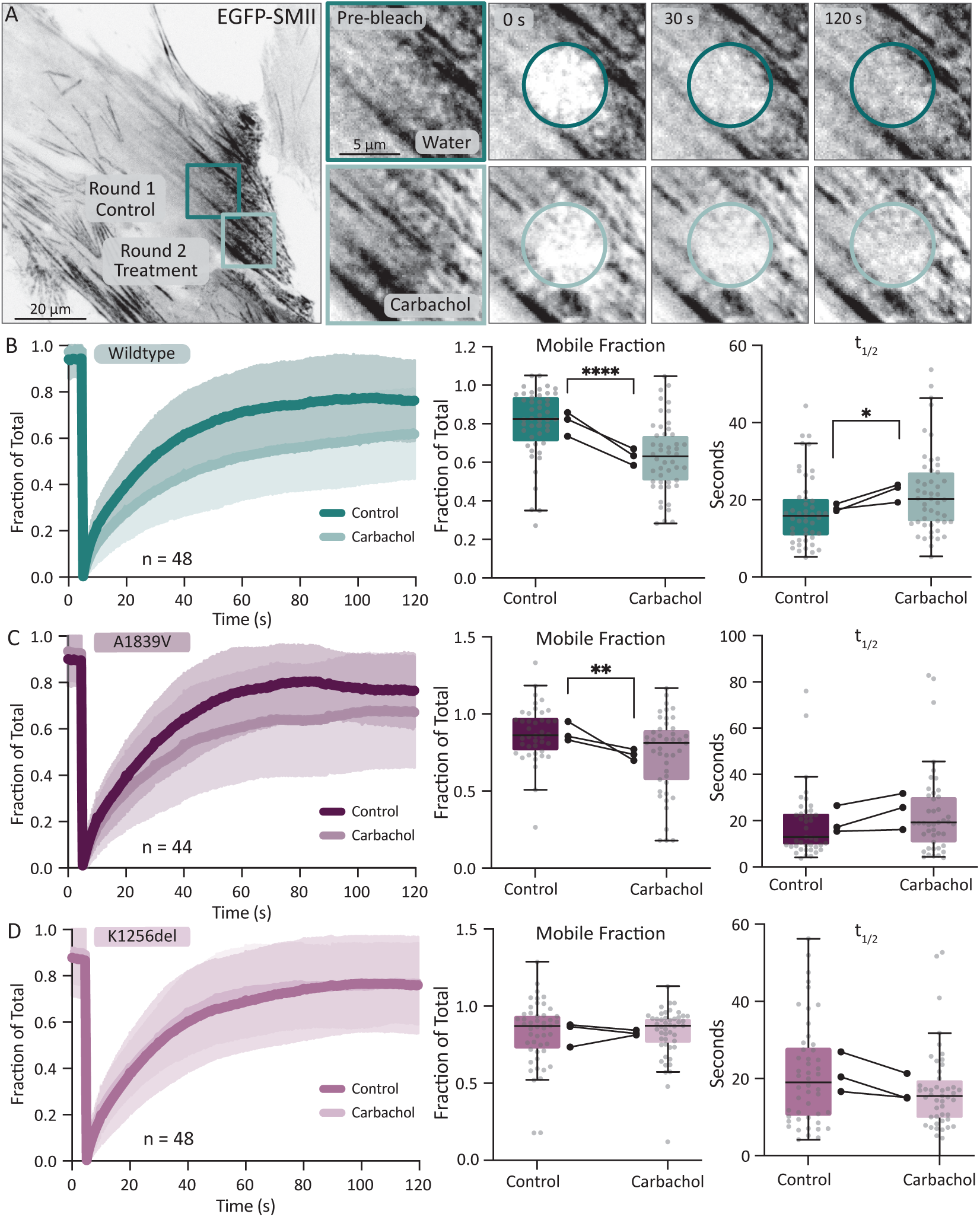
*K1256del abolishes agonist-induced stabilization of SMII filaments.* **A)** Representative paired FRAP experiments in a VSMC expressing EGFP-SMII. FRAP was first performed in one region under control conditions, after which carbachol was added to a final concentration of 10 *μ*mol/L and FRAP was performed 30–60 seconds later in a second, distinct region of the same cell. Circles indicate the bleached regions before bleaching and during fluorescence recovery. **B–D)** Mean FRAP recovery curves and quantification of *t*_1_*_/_*_2_ and mobile fraction before and after treatment with 10 *μ*mol/L carbachol in cells expressing B) wild-type, C) A1839V, or D) K1256del EGFP-SMII. Recovery curves are shown as mean *±* SEM and were fit to a single exponential. Individual-cell data are shown as small gray points with box-and-whisker plots showing the median and interquartile range. Larger black points represent independent experimental-replicate means, with lines connecting matched replicate means between control and carbachol conditions. Wild-type, *n* = 48; A1839V, *n* = 44; K1256del, *n* = 48. Control and carbachol conditions were compared using paired two-sided t tests. * indicates *p <* 0.05; ** indicates *p <* 0.01; **** indicates *p <* 0.0001.

Together, these results demonstrate that *MYH11* tail variants differentially impair the dynamic regulation of SMII filaments. A1839V retained a partial stabilization response to agonist activation, whereas K1256del was dynamically unresponsive to the same stimulus. Because VSMCs must rapidly tune force generation to regulate vascular tone and withstand changing mechanical loads, failure of K1256del SMII to stabilize during activation provides a mechanistic link between altered SMII filament dynamics, reduced contractile output, impaired force-associated adhesion organization, and vulnerability of the arterial wall in *MYH11*-associated aortic disease.

## Discussion

Vascular smooth muscle function depends on a paradox: SMII must be both dynamic and stable [10, 33, 14]. At baseline, SMII filaments undergo molecular exchange, allowing the contractile units and cytoskeleton to remodel. During contractile activation, however, SMII filaments assemble and stabilize to transmit force through the actomyosin cytoskeleton [14]. This regulated transition may be especially vulnerable to disease-associated variants in the *MYH11* tail, which mediates the higher-order assembly of SMII filaments [11]. Using a novel VUS A1839V and the previously defined pathogenic variant K1256del, we show that these *MYH11* mutations disrupt SMII organization and function at the cellular level, providing a tractable approach to categorizing the impact of other VUS.

This study provides, to our knowledge, the first direct visualization of these *MYH11* variants in individual VSMCs. We observed two distinct phenotypes: A1839V caused measurable but milder defects, whereas K1256del broadly disrupted SMII organization and dynamic regulation. Specifically, we found that *MYH11* K1256del reduced SMII incorporation into long stress fibers, increased cytoplasmic and aggregate-like SMII organization, reduced the fraction of assembled SMII, increased SMII network flow, and accelerated SMII exchange kinetics. Combined, these findings demonstrate that *MYH11* K1256del impairs the formation of a stable, force-bearing SMII filament network. This is consistent with prior work showing that *MYH11* K1256del reduces aortic structural integrity and contractility in mice, but extends those observations by defining multiple cell-level mechanisms.

Impaired contractile network formation is relevant to aortic disease progression because the aorta must withstand repetitive pulsatile pressure with every heartbeat [6]. As the dominant cellular component of the medial layer, VSMCs contribute to vascular tone, wall integrity, and the ability of the vessel to adapt to mechanical load through the regulation of SMII activity. If SMII are unstable at baseline or fail to stabilize during activation, the VSMC contractile apparatus may be less able to maintain or adapt tension as needed. Over time, this could weaken the ability of the medial layer to counteract luminal pressure, providing a plausible cellular mechanism by which *MYH11* tail variants such as K1256del predispose the vessel wall to aneurysm growth or dissection.

Our focal adhesion and traction force data further support this model, as VSMC contraction requires more than SMII motor activity alone. Force must be transmitted through actin stress fibers and cell-matrix adhesions to influence tissue mechanics [5, 34]. *MYH11* K1256del-expressing cells showed reduced contractile ability and reduced phospho-paxillin-positive adhesion area, demonstrating that the variant disrupts both components required for force transmission. This is consistent with previous work showing that focal adhesions in VSMCs are mechanically active sites of force transmission and that disruption of focal adhesion components can impair SMC-matrix coupling, disrupt aortic wall organization, and promote aneurysmal remodeling [34, 35, 36]. VSMCs carrying *MYH11* R247C, a pathogenic motor domain variant, displayed altered, less mature focal adhesions together with reduced RhoA activation, further supporting a link between impaired SMII contractility, focal adhesion organization, and downstream mechanosensitive signaling [19]. Because the actin cytoskeleton is also mechanically coupled to the nuclear lamina through the LINC complex, impaired actomyosin tension could additionally alter nuclear mechanotransduction [37].

The disrupted cytoskeletal organization and impaired contractility both point to fundamental issues with myosin filament assembly and function. Myosin 2 filament assembly depends on specific parallel and antiparallel interactions between coiled-coil rods, with charge distributions along the tail helping determine molecular packing [11, 38]. Therefore, deleting a residue along the coiled-coil, as in K1256del, will likely generate a downstream destabilizing mismatch in electrostatic interactions between coiled-coils. K1256 is roughly 40% of the way down the coiled-coil, meaning the remaining 60% in the K1256del will experience disrupted electrostatic interactions. Therefore, this single deletion event could propagate down the entire tail, resulting in the significant assembly and functional defects we observe in our studies. In contrast, A1839V lies near the C-terminus of the coiled-coil in the "assembly competence domain" or ACD, a critical domain for filament assembly. While replacing alanine with valine is a conservative hydrophobic substitution, the bulkier valine in A1839V monomers may sterically disrupt tail-tail interactions during filament assembly, although to a lesser extent than the pathogenic K1256del. Better structural data, especially of SMII tails in filaments, will help clarify these speculative models.

The severity of these assembly defects results in varying functional defects. In our previous work, contractile activation increased wild-type SMII assembly and reduced molecular exchange, consistent with agonist-induced filament stabilization [14]. Here, carbachol again stabilized wild-type EGFP-SMII, reducing mobile fraction and increasing *t*_1_*_/_*_2_. A1839V retained part of this response, showing reduced mobile fraction after stimulation but no increase in *t*_1_*_/_*_2_. In contrast, K1256del failed to respond to carbachol. Neither mobile fraction nor *t*_1_*_/_*_2_ changed significantly, indicating that K1256del abolishes the normal ability of SMII to transition into a more stable filament state during activation, providing a mechanistic link between altered SMII dynamics and impaired sustained force generation under physiologic mechanical demand.

These findings raise the possibility that the impaired contractile function we observed in these tail variants could be pharmacologically rescued by directly increasing myosin activity. In cardiac muscle, small-molecule myosin activators such as omecamtiv mecarbil and danicamtiv increase recruitment of force-generating myosin heads and alter cross-bridge cycling to augment contractility [39, 40, 41]. Although these compounds are selective for cardiac myosin, they establish proof of principle that myosin motor activity can be pharmacologically modulated. Developing or identifying similar SMII-selective activators could therefore provide a strategy to test whether increasing myosin activity can restore force generation in *MYH11* variants in culture. While SMII is found in smooth muscle tissue throughout the body, tissue-specific targeting approaches might enable intervention in diseased or susceptible vasculature.

The A1839V variant presented a milder phenotype than K1256del in our study, with reduced SMII assembly and impaired force generation despite retaining more recognizable filamentous organization. Importantly, A1839V has not been established as a TAAD-causing variant, and the patient carrying A1839V did not have thoracic aortic enlargement. The patient did, however, have severe infrarenal abdominal aortic and bilateral common iliac aneurysmal disease, while A1839V reproducibly altered SMII behavior in our cellular assays. These observations do not establish that A1839V caused the patient’s vascular disease, but they raise the possibility that functionally consequential *MYH11* variants may contribute to a broader spectrum of vascular phenotypes than classic TAAD/PDA or act as lower-penetrance risk alleles.

There is precedent for this possibility. A likely pathogenic *MYH11* p.Arg254Cys variant was identified in a patient with an abdominal aortic aneurysm and segregated to her son, who had bilateral iliac aneurysms and a small abdominal aortic aneurysm [42]. *MYH11* mutations have also been reported in cerebral steno-occlusive arteriopathy and ischemic stroke [43]. These findings suggest that the phenotypic consequences of *MYH11* variation may extend beyond a simple binary classification of TAAD-causing versus benign and may depend on variant-specific penetrance, genetic background, environmental factors, and the vascular tissue involved.

This uncertainty highlights the potential value of cellular phenotyping for *MYH11* VUS. Clinical sequencing continues to identify rare *MYH11* variants, many of which lack sufficient segregation, population, or functional evidence for confidentclassification [44]. At the time of submission, ClinVar listed 1,258 germline *MYH11* variants classified as VUS and a further 144 with conflicting classifications, compared with only 54 classified as pathogenic (*n* = 26) or likely pathogenic (*n* = 28) [45]. VUS thus outnumber established disease-causing variants by more than 20-fold, leaving patients and at-risk relatives without a clear basis for aortic surveillance or cascade testing. Generating mouse models for each variant is not feasible. Cell-based visualization of SMII therefore provides a scalable way to identify variant-specific biological effects and prioritize variants for deeper study. This approach cannot establish clinical pathogenicity or replace genetic and in vivo evidence, but it can provide mechanistic information that is otherwise unavailable for many poorly characterized variants.

It is important to acknowledge and consider limitations of this approach. First, these experiments used adenoviral expression of EGFP-tagged SMII in cultured VSMCs that retain endogenous SMII. Although expression-level analyses suggest that the endogenous SMII is present at low levels, and that the observed phenotypes are not proportionate to EGFP-SMII expression level, this system does not fully reproduce endogenous heterozygous expression of the variants. At the same time, the ability of variant EGFP-SMII to produce clear structural and mechanical defects despite the continued presence of endogenous wild-type SMII is consistent with a dominant-negative effect. This is particularly relevant because *MYH11*-associated aortic disease is typically caused by heterozygous variants, and prior studies of disease-associated *MYH11* coiled-coil variants have demonstrated interactions between mutant and wild-type myosin consistent with dominant-negative behavior [16]. In addition, heterozygous K1256del mice develop aortic dissection following angiotensin II challenge [20]. However, because our adenoviral system expresses variant SMII above endogenous levels, it does not reproduce native heterozygous expression and therefore cannot directly establish a dominant-negative mechanism.

Second, cultured VSMCs do not fully capture the three-dimensional architecture, cyclic strain, matrix composition, and lamellar organization of the aortic media. VSMCs are known to undergo phenotypic modulation in culture, and prior work has shown that impaired *MYH11* function can promote a less differentiated SMC phenotype, including increased proliferation and reduced expression of contractile markers [46, 19, 47]. Cellular phenotyping studies of relevant variants followed by animal models could alleviate these issues.

Together, these findings support a model in which *MYH11* tail variants impair VSMC function by disrupting the dynamic organization, assembly, and stabilization of SMII filaments. K1256del, a known pathogenic variant, prevents formation of a stable, force-bearing SMII network, leading to reduced contractile output, impaired adhesion organization, increased network flow, accelerated exchange, and loss of agonist-induced stabilization. *MYH11* VUS A1839V produces milder but measurable defects despite its unresolved clinical significance. This quantitative molecular data at the single cell level suggest variants like A1839V that mildly alter SMII function likely only lead to disease when secondary genetic or environmental insults are present, while more disruptive variants like K1256del are sufficient on their own to drive disease. More broadly, this work establishes live-cell SMII phenotyping as a practical approach for connecting human *MYH11* variation to cytoskeletal mechanism and for identifying variants that may compromise the mechanical integrity of vascular smooth muscle.

## Materials and Methods

### Clinical identification of the *MYH11* A1839V variant

The *MYH11* A1839V variant was identified in a patient evaluated at Loyola Medicine for aortic aneurysmal disease. The patient had severe infrarenal abdominal aortic and bilateral common iliac aneurysmal disease requiring repeated endovascular repair and a family history notable for abdominal aortic aneurysm in multiple first-degree relatives. Available clinical imaging did not demonstrate thoracic aortic enlargement. Genetic testing was performed on peripheral blood using a clinical multigene hereditary aortopathy panel, which identified the *MYH11* NM_002474.3:c.5516C>T (p.Ala1839Val) variant. The patient provided written informed consent for the use and publication of their clinical and genetic information. This study was approved by the Loyola University Chicago Institutional Review Board committee. Only de-identified clinical information and variant sequence data were used, in accordance with the HIPAA Privacy Rule. The study was conducted in accordance with the principles of the Declaration of Helsinki. No samples or data from relatives were collected. Family history was obtained from the patient’s clinical record.

### MYH11 constructs, VSMC culture, and adenoviral expression

Wild-type, A1839V, and K1256del EGFP-SMII constructs encoded the human SM1A isoform with EGFP fused to the N-terminus of the MYH11 heavy chain. Constructs were generated using the same cloning strategy described previously [14], with the corresponding *MYH11* variant sequence incorporated into the A1839V or K1256del construct. Constructs were validated by whole-plasmid sequencing and packaged into adenoviral vectors by VectorBuilder at titers ranging from 1.01 *×* 10^10^ to 1.14 *×* 10^10^ IFU/mL. Detailed cloning procedures are provided in the SI Appendix.

Primary human aortic VSMCs (ATCC PCS-100-012) were maintained in vascular cell basal medium supplemented with the ATCC Vascular Smooth Muscle Cell Growth Kit and 1% antibiotic–antimycotic. Cells were maintained at 37*^◦^*C and 5% CO_2_, used through passage 4, and tested weekly for mycoplasma. Cells were transduced with wild-type, A1839V, or K1256del EGFP-SMII adenovirus diluted 1:1000 from purified stocks. Medium was replaced after 24 hours, and experiments were performed 48 hours after transduction.

### Imaging and functional assays

Live-cell imaging was performed on a Yokogawa W1 spinning-disk confocal microscope mounted on a Zeiss Axio Observer 7 with a 63*×*/1.4 NA Plan-Apochromat objective. Fixed-cell super-resolution imaging was performed using a Zeiss Elyra 7 with Lattice SIM^2^. Detailed imaging conditions are provided in the SI Appendix.

SMII architecture was classified into five categories: long stress fibers, dense networks, short stress fibers, cytoplasmic, and aggregates, by three reviewers blinded to variant identity. At least 150 cells were evaluated per variant. Focal adhesion organization was assessed by phospho-paxillin immunofluorescence, with total phospho-paxillin-positive adhesion area normalized to whole-cell area.

Traction force microscopy was performed on fibronectin-coupled 8 kPa polyacrylamide gels containing fluorescent microspheres using previously described methods [14]. Bead displacements were calculated using optical flow and traction stresses using Fourier transform traction cytometry. Contractile energy was normalized to cell area. Spatial force organization was additionally assessed by measuring traction stress intensity as a function of distance from the cell edge.

SMII network dynamics were measured from 1-minute interval time-lapse EGFP-SMII images using dense optical flow. Flow-vector magnitude was calculated at each pixel within the cell mask, and mean network flow for each cell was calculated by averaging flow-vector magnitudes across the masked cell area and imaging period. Pixel displacement was converted using the image calibration (0.1746 *μ*m/pixel), with 1 pixel/min corresponding to 0.00291 *μ*m/s.

SMII filament assembly was measured using a single-cell Triton permeabilization assay as described previously [14]. The fraction assembled was calculated as background-subtracted EGFP-SMII signal remaining after permeabilization divided by the pre-permeabilization signal.

FRAP was performed using a 4-*μ*m circular bleach region and fluorescence recovery was measured for 120 seconds. Recovery curves were fit to a single exponential to determine mobile fraction and *t*_1_*_/_*_2_. For agonist experiments, FRAP was first performed under control conditions, followed by addition of carbachol to a final concentration of 10 *μ*mol/L carbachol and a second FRAP measurement in a separate region of the same cell 30–60 seconds later. Full assay and analysis procedures are provided in the SI Appendix.

### Statistical analysis

Statistical analyses were performed using GraphPad Prism version 11.0.2. All tests were two-sided, and *p <* 0.05 was considered statistically significant. Independent experiments were defined as experiments performed on different days using independent cell platings and viral transductions. Unless otherwise indicated, the individual cell was the statistical unit. Numbers of cells and independent experimental replicates are reported in the corresponding figure legends. Small gray points represent individual cells and larger points represent independent experimental-replicate means where shown.

Normality was assessed using the Shapiro-Wilk test. SMII morphology distributions were compared among wild-type, A1839V, and K1256del using a chi-square test of independence on the scorer-normalized 3*×*5 contingency table. Signed Pearson residuals were used to identify variant morphology combinations contributing most strongly to the overall difference, with *|r| >* 2 considered a major contribution.

Focal adhesion area normalized to cell area and contractile energy per area were compared among groups using Kruskal-Wallis tests followed by Dunn’s multiple-comparisons tests comparing A1839V and K1256del with wild-type. Mean SMII network flow was analyzed using planned pairwise Mann-Whitney U tests comparing each variant with wild-type because these distributions were not normally distributed. Baseline SMII assembled fraction, FRAP *t*_1_*_/_*_2_, and FRAP mobile fraction were analyzed using separate unpaired two-sided t tests comparing A1839V and K1256del with wild-type. Paired control and carbachol FRAP measurements within each genotype were analyzed using paired two-sided t tests.

Box-and-whisker plots show the median and interquartile range. Time-course data are presented as mean *±* SEM. Signifi-cance is indicated as \**p <* 0.05, \*\**p <* 0.01, \*\*\**p <* 0.001, and \*\*\*\**p <* 0.0001.

### Data and code availability

Data supporting the findings of this study are available from the corresponding authors upon reasonable request. Custom code used for traction force microscopy, FRAP, and dense optical flow analysis is available at https://github.com/OakesLab.

## Acknowledgements

We thank members of the Beach and Oakes labs for helpful comments and the Loyola University Chicago Core Microscopy Facility. This work was supported in part by National Institutes of Health grants R35-GM138183 to J.R.B, R01-GM148644 to P.W.O., F30-HL178288 to S.K.D., and shared instrument grant S10OD034431 for Zeiss SIM5 to Loyola University Chicago.

## Supplementary Information

### SI Materials and Methods

#### Mammalian expression constructs and adenoviral vectors

The wild-type EGFP-SMII construct encoded the SM1A isoform of human smooth muscle myosin II with EGFP fused to the N-terminus of the MYH11 heavy chain. The parental pEGFP-SM1A construct was generated as described previously [14]. Briefly, a single gBlock purchased from Integrated DNA Technologies contained the 5*^′^* 420 base pairs of human SM1A, including the naturally occurring SalI restriction site, a 5-base-pair linker to facilitate restriction enzyme digestion, and the terminal 3*^′^* 616 base pairs, including the naturally occurring BlpI restriction site and terminal stop codon. This fragment was inserted into pEGFP-C1 following digestion with BglII and KpnI. The internal *MYH11* coding sequence not included in the gBlock was isolated from full-length human *MYH11* cDNA (MHS6278-202857900; Horizon Discovery) by digestion with SalI and BlpI and ligated into the pEGFP-C1-SM1A intermediate. The A1839V and K1256del EGFP-SMII constructs were generated using the same cloning strategy, with the corresponding variant sequence incorporated into the SM1A coding sequence. The A1839V construct contained MYH11 c.5516C>T, resulting in p.Ala1839Val. The K1256del construct contained deletion of one lysine residue at codon 1256. All variant-containing regions and cloning junctions were validated by whole-plasmid sequencing. Wild-type, A1839V, and K1256del EGFP-SMII constructs were packaged into adenoviral vectors by VectorBuilder at titers ranging from 1.01 *×* 10^10^ to 1.14 *×* 10^10^ IFU/mL..

### VSMC CULTURE AND ADENOVIRAL TRANSDUCTION

Primary human aortic VSMCs (ATCC PCS-100-012) were maintained in vascular cell basal medium (ATCC PCS-100-030) supplemented with recombinant human FGF-basic (5 ng/mL), insulin (5 *μ*g/mL), EGF (5 ng/mL), L-glutamine (10 mM), ascorbic acid (50 *μ*g/mL), 5% fetal bovine serum (ATCC Vascular Smooth Muscle Cell Growth Kit, PCS-100-042), and 1% antibiotic–antimycotic solution (MT30004CI; Corning). Cells were maintained at 37*^◦^*C in a humidified incubator containing 5% CO_2_ and used through passage 4. Cells were tested weekly for mycoplasma contamination. Cells were transduced with wild-type, A1839V, or K1256del EGFP-SMII adenovirus at 1:1000. Medium was replaced after 24 hours and experiments were performed 48 hours after transduction.

### Microscopy

Unless otherwise indicated, imaging was performed using a Yokogawa W1 spinning-disk confocal microscope mounted on a Zeiss Axio Observer 7 with a Phasor spatial illumination system and a 63*×*/1.4 NA Plan-Apochromat objective. Live-cell imaging was performed at 37*^◦^*C and 5% CO_2_, with Zeiss Definite Focus used to maintain focal position. Imaging parameters were held constant within each experiment. Images were processed using FIJI/ImageJ, SlideBook 6, and custom Python scripts. Brightness and contrast adjustments were linear and applied consistently across conditions.

For SMII/NM2A analysis, individual filament bundles were cropped from reconstructed images and line scans drawn along individual fibers in FIJI/ImageJ. EGFP-SMII and NM2A fluorescence intensities were background-subtracted and normalized to the maximum signal within each channel for display.

### Western blotting

Cells were scraped and resuspended in 2*×* sample buffer, boiled for 5 minutes, briefly sonicated, and used immediately or stored at *−*20*^◦^*C. Proteins were separated on 7.5% polyacrylamide gels and transferred to PVDF membranes using an eBlot L1 system (GenScript).

Membranes were blocked in 3% BSA in PBST and incubated overnight at 5*^◦^*C with antibodies against SMII heavy chain/MYH11 (A4064; ABclonal; 1:2000), EGFP/GFP (SC-9996; Santa Cruz Biotechnology; 1:2000), and *β* -actin (Gene-Tex 109639; 1:2000). Membranes were incubated with HRP-conjugated secondary antibodies for 1 hour at room temperature. Uncropped blots are provided in (Fig. S4).

### Immunocytochemistry

Cells were fixed with 4% paraformaldehyde in cytoskeletal buffer containing 0.1 M MES, 0.03 M MgCl_2_, 1.38 M KCl, and 0.02 M EGTA for 10 minutes at 37*^◦^*C. Cells were permeabilized with 0.5% Triton X-100 in cytoskeletal buffer for 10 minutes and blocked with 10% goat serum in PBS for 30 minutes. Primary antibodies were diluted in 5% goat serum and incubated overnight at 4*^◦^*C. Antibodies included phospho-paxillin (44-722G; Invitrogen; 1:200) and NM2A/MYH9 (MP3791; ECM Biosciences; 1:200). Cells were washed three times with 0.05% Triton X-100 in PBS and incubated with secondary antibodies for 1 hour at room temperature. Actin was labeled with Alexa Fluor 405-conjugated phalloidin (A30104; Thermo Fisher Scientific; 1:500). Samples were mounted using ProLong Glass antifade mountant (P36980; Thermo Fisher Scientific).

### SMII MORPHOLOGY CLASSIFICATION

Cells were classified into five prespecified SMII morphology categories: long stress fibers, dense networks, short stress fibers, cytoplasmic, and aggregates. The same image set was independently evaluated by three reviewers blinded to variant identity. At least 150 cells were evaluated per variant. Individual cells could be assigned to more than one morphology category. Because total classifications differed among reviewers, classifications for each reviewer and variant were normalized to the total classifications assigned by that reviewer. Scorer-specific fractions were averaged within each variant and multiplied by the mean total number of classifications for that variant to generate a scorer-normalized summary contingency table. Expected counts for the chi-square analysis were calculated from row and column totals. Signed Pearson residuals were calculated as *r* = *^O^√^−E^* , where *O* and *E* represent observed and expected counts, respectively. Positive residuals indicate categories observed more frequently than expected, whereas negative residuals indicate categories observed less frequently than expected. Residuals with an absolute value greater than 2 were considered major contributors to the overall distributional difference and are summarized in Table S1.

### Fixed-cell focal adhesion quantification

Whole-cell masks were generated from the EGFP-SMII channel using thresholding. Phospho-paxillin-positive adhesions were segmented from background-subtracted images using identical processing parameters across variants. Total phospho-paxillin-positive adhesion area and whole-cell area were measured for each cell, and adhesion area was normalized to whole-cell area.

### Traction force microscopy

Traction force microscopy was performed as described previously [14]. Glass coverslips were first activated with 2% 3-aminopropyltrimethoxysilane (313255000; Acros Organics) in isopropanol followed by 1% glutaraldehyde (16360; Electron Microscopy Sciences). Polyacrylamide gels with a shear modulus of approximately 8 kPa were prepared using final concentrations of 7.5% acrylamide and 0.30% bis-acrylamide. Gels contained 0.04-*μ*m fluorescent microspheres (F8789; Invitrogen), were rehydrated overnight, and were coupled to human plasma fibronectin using Sulfo-SANPAH. VSMCs were plated on gels and allowed to adhere overnight. Images of cells and fluorescent beads were acquired before cells were removed using 0.05% SDS to obtain an unstrained reference bead image.

Traction force analysis was performed using custom Python code (https://github.com/OakesLab/TFM). Images were flat-field corrected and aligned to the reference image. Bead displacements were calculated using an OpenCV optical-flow algorithm with an 8-pixel window. Traction stresses were calculated using Fourier transform traction cytometry with a regularization parameter of 2.0 *×* 10*^−^*^6^. Contractile energy was calculated as one-half the summed product of substrate displacement and traction vectors beneath the cell and normalized to cell area derived from the EGFP-SMII image.

### Spatial analysis of traction force organization

Traction maps were classified as containing organized internal force patterns or lacking internal force patterns. Internal forces were defined as traction stress signal persisting into the cell interior rather than being restricted primarily to the cell boundary. Classification was performed manually while blinded to variant identity. For spatial profiling, each pixel within the cell mask was assigned a distance from the nearest cell boundary. Traction stress-map intensity was binned by distance from the cell edge from 0 to 200 pixels. For group profiles, mean traction stress-map intensity was calculated within each distance bin for each cell and then averaged across cells within each variant; shaded regions represent SEM.

### EGFP-SMII expression-intensity analysis

Mean EGFP-SMII fluorescence intensity was measured within each cell mask after local background subtraction. EGFP-SMII intensity was plotted against contractile energy per area or mean SMII network flow to evaluate whether the observed phenotypes were related to expression level.

### Dense optical flow analysis

Time-lapse EGFP-SMII images were acquired at 1-minute intervals for 60 minutes. The image calibration was 0.1746 *μ*m/pixel; therefore, 1 pixel/min corresponded to 0.00291 *μ*m/sec. Image sequences were background-corrected and corrected for photo-bleaching. Cell masks were generated from the EGFP-SMII channel. Dense optical flow was calculated between consecutive frames using the Farneback dense optical flow function in OpenCV with a window size of 48 pixels, 4 iterations, 2 levels, a pyramid scale of 0.5, a polynomial order of 7 and sigma of 1.25. The resultant magnitude of the flow vector was calculated at each pixel and restricted to the cell mask. Mean network flow for each cell was then calculated by averaging these local flow magnitudes across the masked cell area and imaging period.

### Single-cell SMII assembly assay

The Triton permeabilization assay was performed as described previously [14]. VSMCs were plated on glass-bottom dishes (655891; Greiner) and imaged before permeabilization to obtain total EGFP-SMII signal. Cells were permeabilized using buffer containing 5 μM PEG8000, 100 mmol/L PIPES, 150 mmol/L NaCl, 1 mmol/L EGTA, 1 mmol/L MgCl2, 0.5% Triton. The same positions were re-imaged after permeabilization. Cell masks were manually identified in FIJI/ImageJ. Fraction assembled was calculated as background-subtracted post-permeabilization EGFP-SMII signal divided by background-subtracted pre-permeabilization signal.

### Fluorescence recovery after photobleaching

FRAP was performed using a 4 *μ*m-diameter circular region bleached with the 405 nm laser at 100% power for 500 ms. Images were collected before bleaching and for 120 seconds after bleaching at 10-second intervals. Three regions of interest were analyzed: the bleached region, an unbleached control region within the same cell, and background outside the cell. Background-subtracted fluorescence in the bleached region was normalized to the unbleached control. Recovery curves were fit to

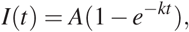

where *A* represents mobile fraction and *k* the observed recovery rate constant. The half-time of recovery was calculated as

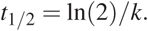

### Carbachol-induced FRAP

A control FRAP experiment was first performed in one region of a VSMC. Carbachol was then added to a final concentration of 10 *μ*M, and FRAP was performed 30–60 seconds later in a separate, previously unbleached region of the same cell. This paired design provided a within-cell control for agonist-induced changes in SMII exchange kinetics.

**Movie M1:** SMII flow in wild-type, A1839V and K1256del mutants associated with Fig. 3A. Flow maps indicate local network direction and velocity. Images were collected every minute for 1 hr and played at a frame rate of 10 fps. Time is in min:sec.

---

**Figure S1.**
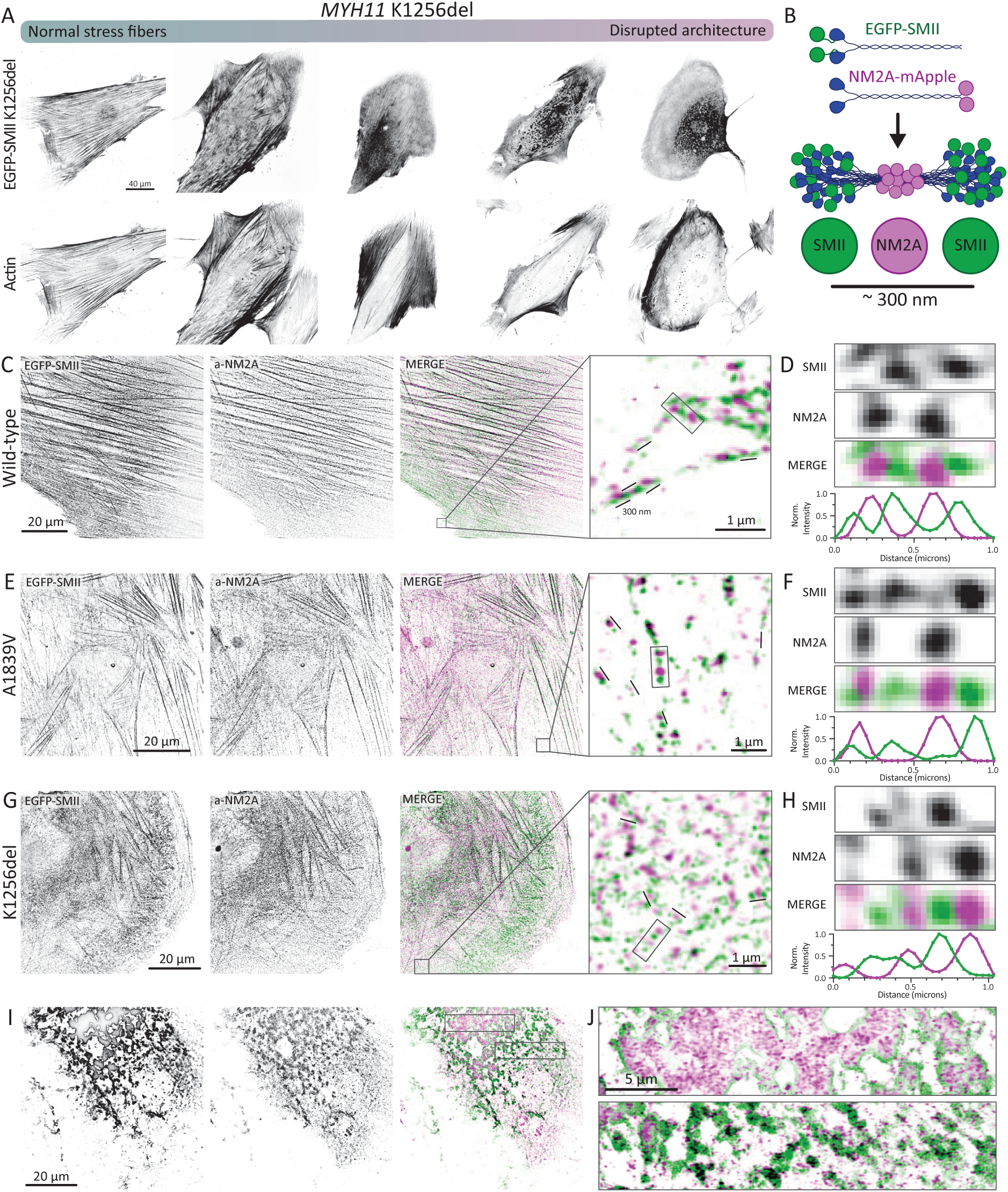
MYH11 tail variants retain the ability to form organized mixed myosin 2 filament structures despite altered higher-order architecture. A) Representative K1256del-expressing VSMCs spanning relatively normal stress fiber organization to severely disrupted SMII architecture. B) Schematic of the organization of EGFP-SMII and NM2A within a mixed myosin 2 filament. C,E,G) Super-resolution images of wild-type C), A1839V E), and K1256del G) EGFP-SMII together with NM2A, with enlarged views of individual filament bundles. D,F,H) Corresponding high-magnification images and line scans showing alternating SMII and NM2A fluorescence patterns in wild-type D), A1839V F), and K1256del H) filament bundles. I,J) Representative K1256del aggregates containing both SMII and NM2A, shown at increasing magnification.

**Figure S2.**
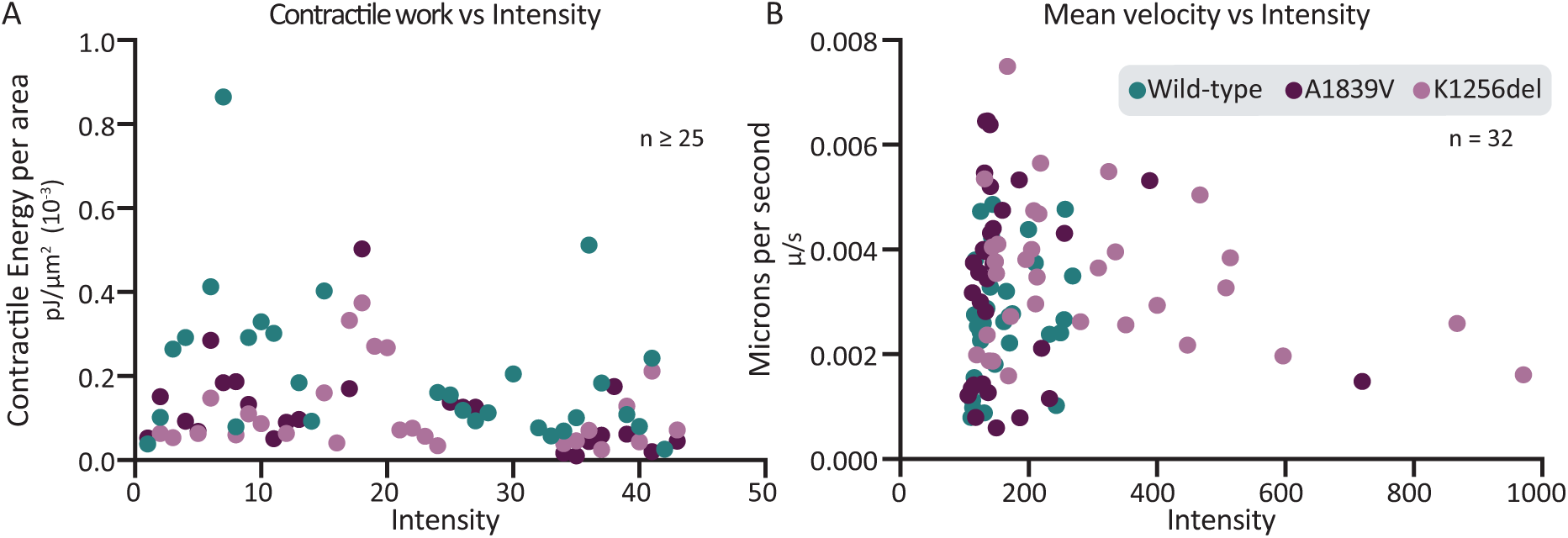
*EGFP-SMII expression level does not explain differences in force generation or SMII network flow.* **A)** Scatter plot of EGFP-SMII fluorescence intensity versus contractile energy per area for cells expressing wild-type, A1839V, or K1256del EGFP-SMII. No obvious relationship was observed between EGFP-SMII expression intensity and traction stress magnitude, suggesting that reduced force generation in variant-expressing cells was not driven simply by differences in EGFP-SMII expression level. **B)** Scatter plot of EGFP-SMII fluorescence intensity versus mean SMII network velocity measured by dense optical flow in cells expressing wild-type, A1839V, or K1256del EGFP-SMII. No obvious relationship was observed between EGFP-SMII expression intensity and mean network velocity, suggesting that increased SMII network flow in K1256del cells was not driven simply by differences in EGFP-SMII expression level.

**Figure S3.**
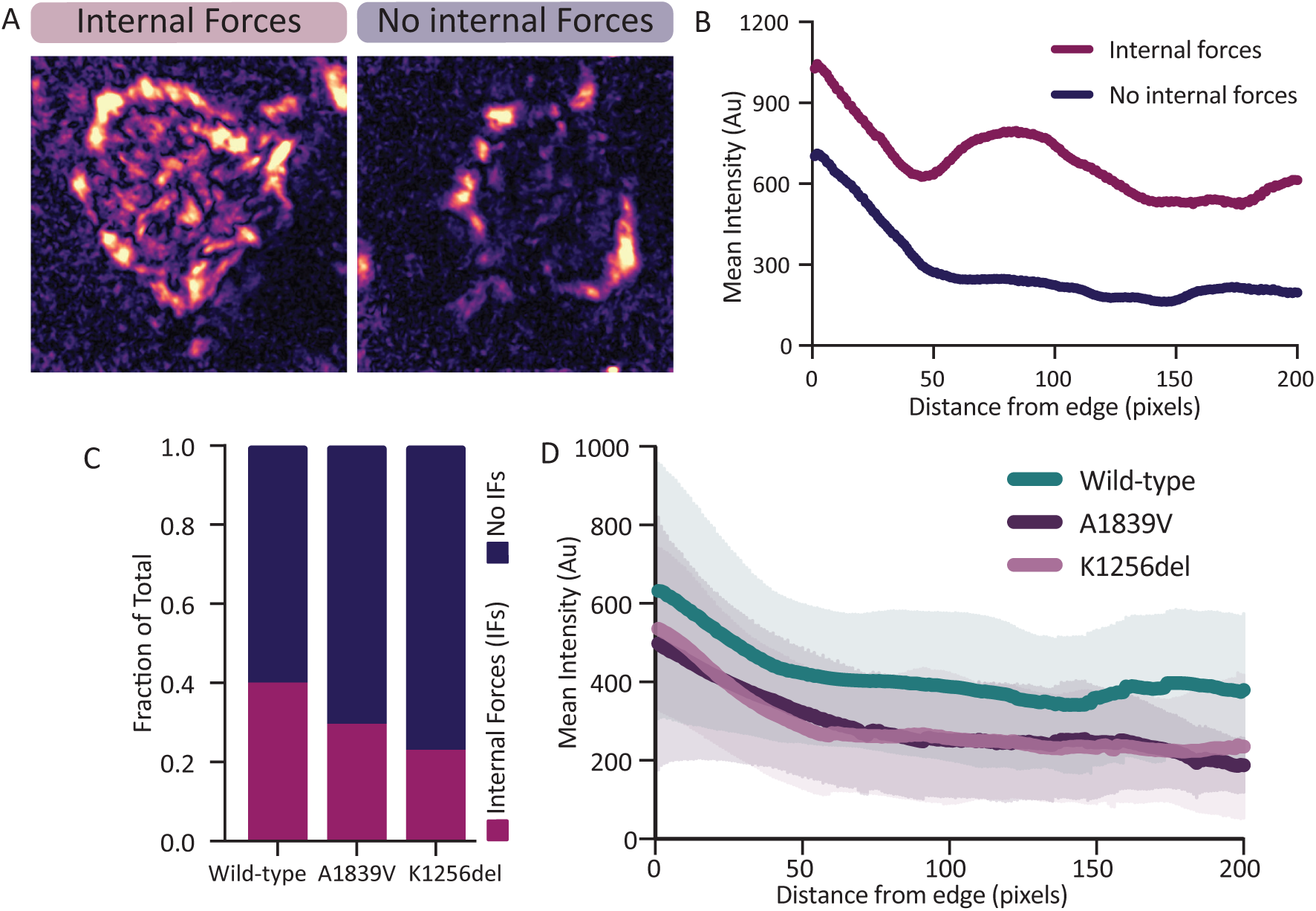
*MYH11 tail variants alter the spatial organization of traction forces.* A) Representative traction stress maps of wild-type EGFP-SMII classified as containing internal forces or lacking internal forces. B) Mean traction stress-map intensity plotted as a function of distance from the cell edge for representative cells with internal forces or no internal forces. Traction maps with internal forces show persistent traction stress signal extending into the cell interior, whereas maps lacking internal forces show a more rapid decline in signal with distance from the edge. C) Quantification of the fraction of cells classified as having internal forces or no internal forces for wild-type, A1839V, and K1256del EGFP-SMII. D) Mean traction stress-map intensity as a function of distance from the cell edge for cells expressing wild-type, A1839V, or K1256del EGFP-SMII. Lines show the group mean, and shaded regions show SEM.

**Figure S4.**
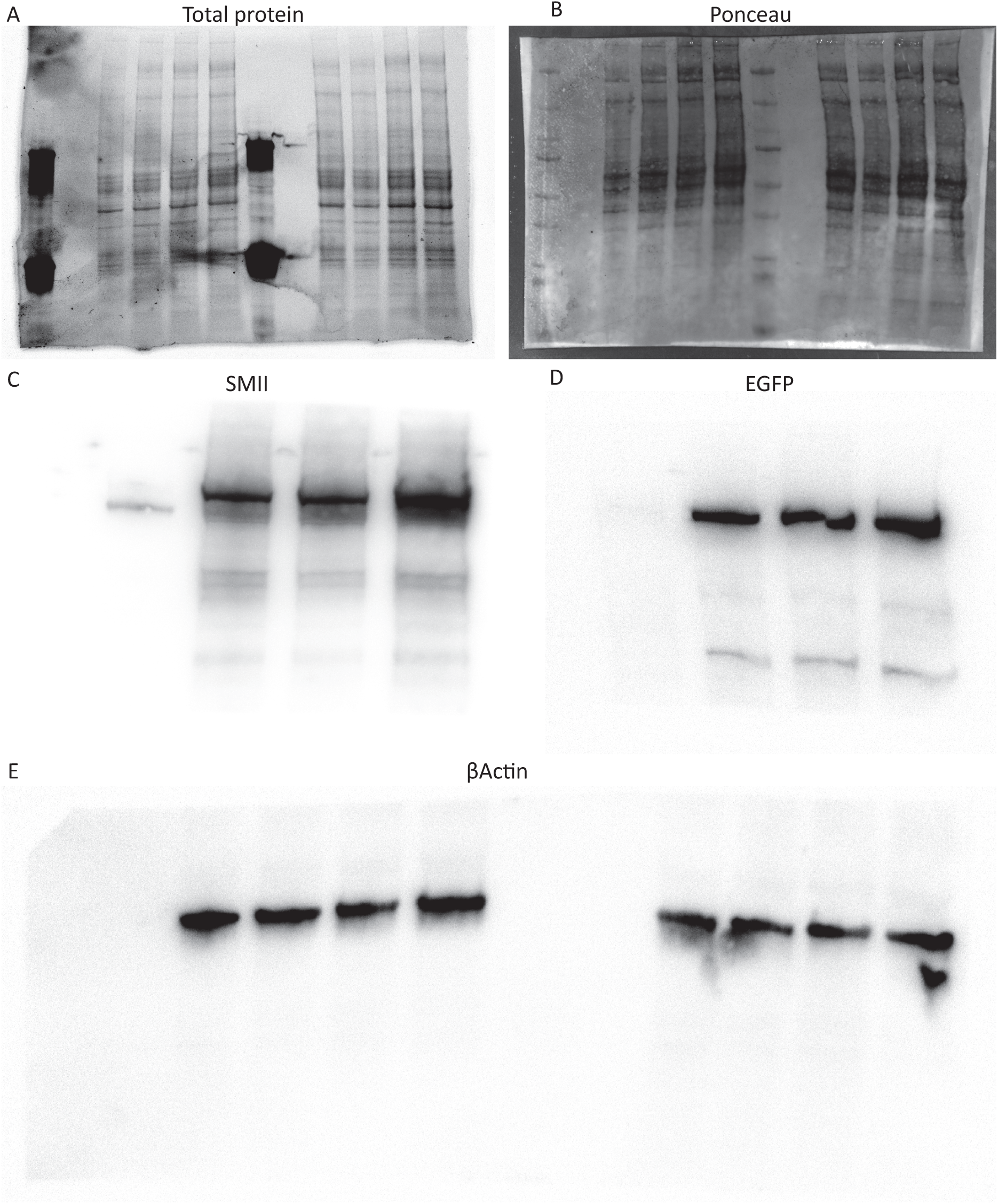
*Full western blots corresponding to* Figure 1B. **A)** Full total-protein image. **B)** Full Ponceau-stained membrane. **C–E)** Full uncropped western blots probed for **C)** SMII/MYH11, **D)** EGFP, and **E)** *β* -actin. The corresponding cropped regions are shown in Figure 1B.

**Table S1.** *MYH11 variant-morphology combinations contributing most significantly to altered SMII architecture.* Signed Pearson residuals from the scorer-normalized 3 *×* 5 chi-square contingency table are shown for variant–morphology combinations with absolute residuals *>* 2. Positive residuals indicate morphologies observed more frequently than expected, whereas negative residuals indicate morphologies observed less frequently than expected.

| Variant | SMII morphology | Residual |
| --- | --- | --- |
| WT | Long stress fibers | + 2.93 |
| WT | Aggregates | - 2.28 |
| A1839V | Short stress fibers | + 2.68 |
| A1839V | Aggregates | - 3.26 |
| K1256del | Aggregates | + 5.51 |
| K1256del | Cytoplasmic | + 2.89 |
| K1256del | Long stress fibers | - 4.80 |

## References

1. G. K. Owens, M. S. Kumar, and B. R. Wamhoff. Molecular regulation of vascular smooth muscle cell differentiation in development and disease. Physiological Reviews, 84(3):767–801, July 2004.

2. R. Yamin and K. G. Morgan. Deciphering actin cytoskeletal function in the contractile vascular smooth muscle cell. The Journal of Physiology, 590(17):4145–4154, September 2012.

3. X. Wang, D. Gonzalez-Rodriguez, T. Vourc’h, P. Silberzan, and A. I. Barakat. Contractility-Induced Self-Organization of Smooth Muscle Cells: From Multilayer Cell Sheets to Dynamic Three-Dimensional Clusters. Communications Biology, 6(1):1–10, 2023.

4. E. C. Davis. Smooth muscle cell to elastic lamina connections in developing mouse aorta. Role in aortic medial organization. Laboratory Investigation; a Journal of Technical Methods and Pathology, 68(1):89–99, January 1993.

5. A. Karimi and D. M. Milewicz. Structure of the Elastin-Contractile Units in the Thoracic Aorta and How Genes That Cause Thoracic Aortic Aneurysms and Dissections Disrupt This Structure. The Canadian Journal of Cardiology, 32(1):26–34, January 2016.

6. J. Humphrey, M. Schwartz, G. Tellides, and D. Milewicz. Role of Mechanotransduction in Vascular Biology: Focus on Thoracic Aortic Aneurysms and Dissections. Circulation research, 116(8):1448–1461, April 2015.

7. D. M. Milewicz, K. M. Trybus, D.-C. Guo, H. L. Sweeney, E. Regalado, K. Kamm, and J. T. Stull. Altered Smooth Muscle Cell Force Generation as a Driver of Thoracic Aortic Aneurysms and Dissections. *Arteriosclerosis*, Thrombosis, and Vascular Biology, 37(1):26–34, 2017.

8. G. J. Ye, A. P. Nesmith, and K. K. Parker. The Role of Mechanotransduction on Vascular Smooth Muscle Myocytes Cytoskeleton and Contractile Function. Anatomical record (Hoboken, N.J. : 2007), 297(9):1758–1769, 2014.

9. I. Y. Kuo and B. E. Ehrlich. Signaling in Muscle Contraction. Cold Spring Harbor Perspectives in Biology, 7(2):a006023, 2015.

10. L. Wang, P. Chitano, and C. Y. Seow. Filament evanescence of myosin II and smooth muscle function. The Journal of General Physiology, 153(3):e202012781, February 2021.

11. R. Straussman, J. M. Squire, A. Ben-Ya’acov, and S. Ravid. Skip residues and charge interactions in myosin II coiled-coils: implications for molecular packing. Journal of Molecular Biology, 353(3):613–628, October 2005.

12. I. M. Tolić-Nørrelykke, J. P. Butler, J. Chen, N. Wang, and New Collective Author. Spatial and temporal traction response in human airway smooth muscle cells. American Journal of Physiology. Cell Physiology, 283(4):C1254–1266, October 2002.

13. S. J. Gunst and W. Zhang. Actin cytoskeletal dynamics in smooth muscle: a new paradigm for the regulation of smooth muscle contraction. American Journal of Physiology -Cell Physiology, 295(3):C576–C587, September 2008.

14. S. K. Demeulenaere, M. A. Bennett, B. Somerfield, H. Wu, M. E. Utgaard, H. Patel, S. Sala, E. R. Longtine, A. Zied, J. A. Kirk, P. W. Oakes, and J. R. Beach. Vascular Smooth Muscle Myosin 2 Filaments Dynamically Assemble and Stabilize During Induced Contractility, April 2026. ISSN: 2692-8205 Pages: 2022.10.10.511341 Section: New Results.

15. A. T. Yetman, L. J. Starr, S. B. Bleyl, L. Meyers, and J. W. Delaney. Progressive Aortic Dilation Associated With ACTA2 Mutations Presenting in Infancy. Pediatrics, 136(1):e262–266, July 2015.

16. L. Zhu, R. Vranckx, P. Khau Van Kien, A. Lalande, N. Boisset, F. Mathieu, M. Wegman, L. Glancy, J.-M. Gasc, F. Brunotte, P. Bruneval, J.-E. Wolf, J.-B. Michel, and X. Jeunemaitre. Mutations in myosin heavy chain 11 cause a syndrome associating thoracic aortic aneurysm/aortic dissection and patent ductus arteriosus. Nature Genetics, 38(3):343–349, March 2006.

17. H. Pannu, V. Tran-Fadulu, C. L. Papke, S. Scherer, Y. Liu, C. Presley, D. Guo, A. L. Estrera, H. J. Safi, A. R. Brasier, G. W. Vick, A. J. Marian, C. S. Raman, L. M. Buja, and D. M. Milewicz. MYH11 Mutations Result in a Distinct Vascular Pathology Driven by Insulin-like Growth Factor 1 and Angiotensin II. Human Molecular Genetics, 16(20):2453–2462, 2007.

18. D. M. Milewicz, D.-C. Guo, V. Tran-Fadulu, A. L. Lafont, C. L. Papke, S. Inamoto, C. S. Kwartler, and H. Pannu. Genetic Basis of Thoracic Aortic Aneurysms and Dissections: Focus on Smooth Muscle Cell Contractile Dysfunction. Annual Review of Genomics and Human Genetics, 9:283–302, 2008.

19. S.-Q. Kuang, C. S. Kwartler, K. L. Byanova, J. Pham, L. Gong, S. K. Prakash, J. Huang, K. E. Kamm, J. T. Stull, H. L. Sweeney, and D. M. Milewicz. Rare, nonsynonymous variant in the smooth muscle-specific isoform of myosin heavy chain, MYH11, R247C, alters force generation in the aorta and phenotype of smooth muscle cells. Circulation Research, 110(11):1411–1422, May 2012.

20. K. Negishi, K. Aizawa, T. Shindo, T. Suzuki, T. Sakurai, Y. Saito, T. Miyakawa, M. Tanokura, Y. Kataoka, M. Maeda, S. Tomida, H. Morita, N. Takeda, I. Komuro, K. Kario, R. Nagai, and Y. Imai. An Myh11 single lysine deletion causes aortic dissection by reducing aortic structural integrity and contractility. Scientific Reports, 12(1):8844, May 2022.

21. Y. Imai, H. Morita, N. Takeda, F. Miya, H. Hyodo, D. Fujita, T. Tajima, T. Tsunoda, R. Nagai, M. Kubo, and I. Komuro. A deletion mutation in myosin heavy chain 11 causing familial thoracic aortic dissection in two Japanese pedigrees. International Journal of Cardiology, 195:290–292, September 2015.

22. M. Kawana, J. A. Spudich, and K. M. Ruppel. Hypertrophic cardiomyopathy: Mutations to mechanisms to therapies. Frontiers in Physiology, 13:975076, 2022.

23. I. Olivotto, A. Oreziak, R. Barriales-Villa, T. P. Abraham, A. Masri, P. Garcia-Pavia, S. Saberi, N. K. Lakdawala, M. T. Wheeler, A. Owens, M. Kubanek, W. Wojakowski, M. K. Jensen, J. Gimeno-Blanes, K. Afshar, J. Myers, S. M. Hegde, S. D. Solomon, A. J. Sehnert, D. Zhang, W. Li, M. Bhattacharya, J. M. Edelberg, C. B. Waldman, S. J. Lester, A. Wang, C. Y. Ho, D. Jacoby, and EXPLORER-HCM study investigators. Mavacamten for treatment of symptomatic obstructive hypertrophic cardiomyopathy (EXPLORER-HCM): a randomised, double-blind, placebo-controlled, phase 3 trial. Lancet, 396(10253):759–769, September 2020.

24. C. A. Miller, A. Quinones-Hinojosa, and S. S. Rosenfeld. Non-muscle myosin II is a promising therapeutic target. Trends in Pharmacological Sciences, 46(10):931–934, October 2025.

25. L. Radnai, E. J. Young, C. Kikuti, K. Toth, M. Zhou, M. Hafenbreidel, R. F. Stremel, L. Lin, P. Pasetto, X. Jin, A. Patel, M. Conlon, S. B. Briggs, L. Heidsieck, H. L. Sweeney, J. Sellers, T. Krieger-Burke, W. H. Martin, J. Sisco, S. Young, P. Pearson, G. Rumbaugh, G. L. Araldi, S. K. Duddy, M. D. Cameron, M. Surman, A. Houdusse, P. R. Griffin, T. M. Kamenecka, and C. A. Miller. Development of clinically viable non-muscle myosin II small molecule inhibitors. Cell, 188(17):4604–4621.e15, August 2025.

26. C. S. Kwartler, L. Gong, J. Chen, S. Wang, R. Kulmacz, X.-y. Duan, A. Janda, J. Huang, K. E. Kamm, J. T. Stull, D. Guo, and D. M. Milewicz. Variants of Unknown Significance in Genes Associated with Heritable Thoracic Aortic Disease Can Be Low Penetrant “Risk Variants”. American Journal of Human Genetics, 103(1):138–143, July 2018.

27. J. C. Sapp, J. J. Johnston, K. Driscoll, A. R. Heidlebaugh, A. Miren Sagardia, D. N. Dogbe, K. L. Umstead, E. Turbitt, I. Alevizos, J. Baron, C. Bönnemann, B. Brooks, S. Donkervoort, Y. H. Jee, W. M. Linehan, F. J. McMahon, J. Moss, J. C. Mullikin, D. Nielsen, E. Pelayo, A. T. Remaley, R. Siegel, H. Su, C. Zarate, NISC Comparative Sequencing Program, T. A. Manolio, B. B. Biesecker, and L. G. Biesecker. Evaluation of Recipients of Positive and Negative Secondary Findings Evaluations in a Hybrid CLIA-Research Sequencing Pilot. American Journal of Human Genetics, 103(3):358–366, September 2018.

28. National Center for Biotechnology Information. ClinVar; VCV000197069.56. https://www.ncbi.nlm.nih.gov/ clinvar/variation/VCV000197069.56. Accessed September 18, 2026.

29. J. R. Beach, L. Shao, K. Remmert, D. Li, E. Betzig, and J. A. Hammer. Nonmuscle myosin II isoforms coassemble in living cells. Current biology: CB, 24(10):1160–1166, May 2014.

30. M. A. Quintanilla, J. A. Hammer, and J. R. Beach. Non-Muscle Myosin 2 at a Glance. Journal of Cell Science, 136(5):jcs260890, 2023.

31. Z. Wang, F. M. Pavalko, and S. J. Gunst. Tyrosine phosphorylation of the dense plaque protein paxillin is regulated during smooth muscle contraction. The American Journal of Physiology, 271(5 Pt 1):C1594–1602, November 1996.

32. H. Satoh. Modulation of Ca(2+)-activated K+ current by isoprenaline, carbachol, and phorbol ester in cultured (and fresh) rat aortic vascular smooth muscle cells. General Pharmacology, 27(2):319–324, March 1996.

33. P. Chitano, L. Wang, G. Y. Y. Tin, M. Ikebe, P. D. Paré, and C. Y. Seow. Smooth muscle function and myosin polymerization. Journal of Cell Science, 130(15):2468–2480, August 2017.

34. Z. Sun, L. A. Martinez-Lemus, M. A. Hill, and G. A. Meininger. Extracellular matrix-specific focal adhesions in vascular smooth muscle produce mechanically active adhesion sites. American Journal of Physiology. Cell Physiology, 295(1):C268–C278, July 2008.

35. D. Shen, J. Li, J. J. Lepore, T. J. Anderson, S. Sinha, A. Y. Lin, L. Cheng, E. D. Cohen, J. D. Roberts, S. Dedhar, M. S. Parmacek, and R. E. Gerszten. Aortic Aneurysm Generation in Mice with Targeted Deletion of Integrin Linked Kinase (ILK) in Vascular Smooth Muscle Cells. Circulation Research, 109(6):616–628, September 2011.

36. Z. Zhu, M. Liu, J. Wei, D. Suryanarayan, P. Behzadi, R. Edgar, J. A. Phillippi, C. S. Hilaire, C. Espinosa-Diez, and D. Gomez. Focal adhesion proteins confer smooth muscle anoikis resistance and protection against aortic aneurysm and dissection. JCI Insight, 11(9), May 2026.

37. L. Porter, R.-M. Minaisah, S. Ahmed, S. Ali, R. Norton, Q. Zhang, E. Ferraro, C. Molenaar, M. Holt, S. Cox, S. Fountain, C. Shanahan, and D. Warren. SUN1/2 Are Essential for RhoA/ROCK-Regulated Actomyosin Activity in Isolated Vascular Smooth Muscle Cells. Cells, 9(1):132, 2020. PMID: 31935926.

38. R. A. Cross, M. A. Geeves, and J. Kendrick-Jones. A nucleation–elongation mechanism for the self-assembly of side polar sheets of smooth muscle myosin. The EMBO Journal, 10(4):747–756, 1991. PMID: 2009856.

39. F. I. Malik, J. J. Hartman, K. A. Elias, B. P. Morgan, H. Rodriguez, K. Brejc, R. L. Anderson, S. H. Sueoka, K. H. Lee, J. T. Finer, R. Sakowicz, R. Baliga, D. R. Cox, M. Garard, G. Godinez, R. Kawas, E. Kraynack, D. Lenzi, P. P. Lu, A. Muci, C. Niu, X. Qian, D. W. Pierce, M. Pokrovskii, I. Suehiro, S. Sylvester, T. Tochimoto, C. Valdez, W. Wang, T. Katori, D. A. Kass, Y.-T. Shen, S. F. Vatner, and D. J. Morgans. Cardiac myosin activation: a potential therapeutic approach for systolic heart failure. Science, 331(6023):1439–1443, March 2011.

40. V. J. Planelles-Herrero, J. J. Hartman, J. Robert-Paganin, F. I. Malik, and A. Houdusse. Mechanistic and structural basis for activation of cardiac myosin force production by omecamtiv mecarbil. Nature Communications, 8(1):190, August 2017.

41. K. B. Kooiker, S. Mohran, K. L. Turner, W. Ma, A. Martinson, G. Flint, L. Qi, C. Gao, Y. Zheng, T. S. McMillen, C. Mandrycky, M. Mahoney-Schaefer, J. C. Freeman, E. G. Costales Arenas, A.-Y. Tu, T. C. Irving, M. A. Geeves, B. C. W. Tanner, M. Regnier, J. Davis, and F. Moussavi-Harami. Danicamtiv Increases Myosin Recruitment and Alters Cross-Bridge Cycling in Cardiac Muscle. Circulation Research, 133(5):430–443, August 2023.

42. K. M. van de Luijtgaarden, D. Heijsman, A. Maugeri, M. M. Weiss, H. J. M. Verhagen, A. IJpma, H. T. Brüggenwirth, and D. Majoor-Krakauer. First genetic analysis of aneurysm genes in familial and sporadic abdominal aortic aneurysm. Human Genetics, 134(8):881–893, 2015. PMID: 26017485.

43. A. Raghuram, S. Sanchez, Y. Lu, M. Hickerson, M. B. Solis Mayorga, J. M. Romero, S. Matsumoto, P. L. Musolino, and E. A. Samaniego. Cerebral arteriopathy and ischemic stroke in a pediatric MYH11 patient. Journal of Stroke and Cerebrovascular Diseases, 32(3):106938, 2023. PMID: 36621119.

44. S. Richards, N. Aziz, S. Bale, D. Bick, S. Das, J. Gastier-Foster, W. W. Grody, M. Hegde, E. Lyon, E. Spector, K. Voelkerding, H. L. Rehm, and ACMG Laboratory Quality Assurance Committee. Standards and guidelines for the interpretation of sequence variants: a joint consensus recommendation of the American College of Medical Genetics and Genomics and the Association for Molecular Pathology. Genetics in Medicine: Official Journal of the American College of Medical Genetics, 17(5):405–424, May 2015.

45. M. J. Landrum, S. Chitipiralla, G. R. Brown, C. Chen, B. Gu, J. Hart, D. Hoffman, W. Jang, K. Kaur, C. Liu, V. Lyoshin, Z. Maddipatla, R. Maiti, J. Mitchell, N. O’Leary, G. R. Riley, W. Shi, G. Zhou, V. Schneider, D. Maglott, J. B. Holmes, and B. L. Kattman. ClinVar: improvements to accessing data. Nucleic Acids Research, 48(D1):D835–D844, January 2020.

46. J. A. Beamish, P. He, K. Kottke-Marchant, and R. E. Marchant. Molecular Regulation of Contractile Smooth Muscle Cell Phenotype: Implications for Vascular Tissue Engineering. Tissue Engineering Part B: Reviews, 16(5):467–491, 2010.

47. J. Burger, N. Bogunovic, N. P. de Wagenaar, H. Liu, N. van Vliet, A. IJpma, A. Maugeri, D. Micha, H. J. M. Verhagen, T. L. M. Ten Hagen, D. Majoor-Krakauer, I. van der Pluijm, J. Essers, and K. K. Yeung. Molecular phenotyping and functional assessment of smooth muscle-like cells with pathogenic variants in aneurysm genes ACTA2, MYH11, SMAD3 and FBN1. Human Molecular Genetics, 30(23):2286–2299, November 2021.

